# Knockdown of PR-DUB subunit *calypso* in the developing *Drosophila* eye and wing results in mis-patterned tissues with altered size and shape

**DOI:** 10.1101/2025.01.09.631961

**Authors:** Max Luf, Priya Begani, Anne M. Bowcock, Cathie M. Pfleger

**Affiliations:** Department of Oncological Sciences, The Icahn School of Medicine at Mount Sinai, New York, New York 10029; The Tisch Cancer Institute, The Icahn School of Medicine at Mount Sinai, New York, New York 10029; The Graduate School of Biomedical Sciences, The Icahn School of Medicine at Mount Sinai, New York, New York 10029

**Keywords:** /phrases: BAP1, Calypso, Polycomb Repression, PR-DUB, *Drosophila*

## Abstract

The deubiquitinating enzyme BAP1, the catalytic subunit of the PR-DUB complex, is implicated in several cancers, in the familial cancer syndrome BAP1 Tumor Predisposition Syndrome, and in the neurodevelopmental disorder Küry -Isidor syndrome. In *Drosophila,* there are numerous reports in the literature describing developmental patterning phenotypes for several chromatin regulators including the discovery of Polycomb itself, but corresponding adult morphological phenotypes caused by developmental dysregulation of *Drosophila BAP1* ortholog *calypso* (*caly*) are less well-described. We report here that knockdown of *caly* in the eye and wing produce concomitant chromatin dysregulation phenotypes. RNAi to *caly* in the early eye reduces survival and leads to changes in eye size and shape including eye outgrowths, some of which resemble homeotic transformations whereas others resemble tumor-like outgrowths seen in other fly cancer models. Mosaic eyes containing *caly* loss-of-function tissue phenocopy *caly* RNAi. Knocking down *caly* across the wing disrupts wing shape and patterning including effects on wing vein pattern. This phenotypic characterization reinforces the growing body of literature detailing developmental mis-patterning driven by chromatin dysregulation and serves as a baseline for future mechanistic studies to understand the role of BAP1 in development and disease.

**ARTICLE SUMMARY:** PR-DUB catalytic subunit deubiquitinating enzyme BAP1 plays an important role in tumor suppression and chromatin regulation. Whereas many chromatin regulators are well-characterized for their roles in patterning, the mis-patterning phenotypes in adult structure for dysregulating BAP1 ortholog *calypso* (*caly*) in development are less well described. We report mis-patterned adult eye and wing phenotypes caused by *caly* RNAi in the developing eye and wing respectively.

## INTRODUCTION

The deubiquitinating enzyme (DUB) BAP1 is a major tumor driver and metastasis suppressor in Uveal Melanoma (UM), the most common primary cancer in the eye [Jager et al., 2022; Kashyap et al., 2016]. *BAP1* is mutated in 45% of UM and in 85% of UM that metastasize [Harbour et al., 2010; Robertson et al., 2017; Field et al., 2018]. In addition, various other cancers have significant loss of *BAP1*, including mesothelioma [Bott et al., 2011; Testa et al., 2011; Wiesner et al., 2011], clear cell renal cancer [Ricketts et al., 2018] and cholangiocarcinoma [Jiao et al., 2013]. Germline mutations in *BAP1* also lead to a *BAP1* Tumor Predisposition Syndrome (BAP1-TPS) [Bergman et al., 2006; Carbone et al., 2012; Carbone et al., 2015]. BAP1 is reported to regulate cell proliferation [Machida et al., 2009], cell death [Bononi et al., 2017], and nuclear processes crucial for genome stability, such as DNA repair and replication [Yu et al., 2014]. The Polycomb repressive system is composed of three main protein complexes: Polycomb Repressive Complex 1 (PRC1), Polycomb Repressive Complex 2 (PRC2), and the Polycomb Repressive Deubiquitinase (PR-DUB) complex, in which BAP1 is the catalytic subunit [Scheuermann et al., 2010]. These complexes repress homeotic (HOX) and other developmental regulator genes in cells where they must stay inactive, thereby ensuring proper differentiation and cellular identity throughout development [Scheuermann et al., 2010]. Loss of essential components of these complexes can result in dysregulation of chromatin organization which leads to developmental abnormalities and diseases such as cancer. Within the PR-DUB complex, BAP1 globally deubiquitinates lysine 119 on histone H2A, and its loss leads to pervasive H2AK119ub1. Failure to constrain pervasive H2AK119ub1 titrates away Polycomb Repressive complexes (PRC) from their targets, decreasing promoter H3K27me3 concentration and also causes chromatin compaction [Daou et al., 2015; Fursova et al., 2021; Conway et al., 2021]. Despite these insights, the specific role of loss of BAP1 function in the development of different cancers and their metastases is unclear.

Model systems such as *Drosophila* provide a useful context to further study BAP1 function and its roles *in vivo*. In fact, *polycomb* was originally discovered in *Drosophila* [Lewis & Mislove, 1947]*. BAP1* is highly conserved and represented by *calypso* (*caly*, also called *dBap1*) in *Drosophila.* Extensive literature in *Drosophila* examines the role of PRC1, PRC2, and a number of Polycomb regulators, but less work has been done to characterize *caly* developmental phenotypes. Previous work in flies showed that loss of *caly* catalytic activity increases monoubiquitinated H2A and decreases repression of PcG target genes and *HOX* genes, as seen in human cell lines and mouse work [Scheuermann et al., 2010; de Ayala Alonso et al., 2007].

Given the importance of *BAP1* in suppression of tumorigenesis and metastasis, it is important to complement the published literature with a detailed description of how *caly* knockdown in development affects patterning and gross morphology across different tissue contexts. This will enable us to further understand the role of *caly* in development and prioritize contexts for further study to elucidate the role *BAP1* as a tumor suppressor in human cancer.

Here we characterize phenotypes arising from knockdown of *caly* using inducible RNAi allele *caly^HMC04109^* in developing tissues. RNAi to *caly* in multiple contexts in the developing eye and wing increased lethality and led to a spectrum of phenotypes in surviving flies. RNAi in the early eye resulted in a range of reduced eye sizes and outgrowths some of which differentiated into structures that resembled antennae, maxillary palps, or even legs. Other outgrowths were difficult to classify and resembled tumor-like outgrowths seen in other fly cancer models. In contrast, RNAi in the differentiating eye results in eyes of slightly increased size that appeared normal morphologically. RNAi in the wing led to abnormalities affecting patterning including wing vein phenotypes, blisters, and crumpling. These phenotypes are similar to those reported due to loss of function of other chromatin regulators and validate the allele *caly^HMC04109^* as a tool in future mechanistic studies to understand the role of BAP1 in disease and cancer.

## RESULTS AND DISCUSSION

Given the importance of BAP1 and the range of phenotypes reported for other polycomb regulators in *Drosophila*, we utilized several Gal4 drivers that drive expression in the developing eye or wing to establish the phenotypes of RNAi allele *caly^HMC04109^*.

### RNAi to or mutation in *caly* in the early eye caused lethality and a range of phenotypes

Driver *ey-gal4* has been characterized to express in the early cells of the imaginal eye disc [Halder et al., 1998; Hazelett et al., 1998; Quiring et al., 1994] and also in other tissues including the nervous system, larval brain, and genital discs [Adachi et al., 2003; Weasner et al., 2009]. Inducing RNAi with *caly^HMC04109^* and *ey-gal4* resulted in substantial pupal lethality compared to controls (quantified in Fig. 1A). *ey>caly^HMC04109^*flies that survived to adulthood demonstrated a range of phenotypes (Fig. 1B-1K) including eyes that appeared “mostly normal” as seen in a previous report [Brown et al., 2023] and rough eyes of reduced size (Fig. 1C-1C’ for males, 1E-1E’ for females; quantified in Fig. 1H; additional examples in Supplemental Fig. S1) compared to control eyes (1B-1B’ for males, 1D-1D’ for females). The heads were reduced in height compared to controls (for anterior view of the eyes, Fig. 1C’ compared to 1B’ for males, 1E’ to 1D’ for females; quantified in Fig. 1I). In many cases, the eye tissue appeared to be bulging (1C’, 1E’), and despite the reduced area at the base of the eye (Fig. 1H), overall width of the head increased compared to controls (Fig. 1J). This is highlighted by an increase in the ratio of width to height compared to control eyes (Supplemental Fig. S2B). Most eyes showed bristle phenotypes in the anterior region in the periphery of the eye (arrows in Fig. 1C), and we also saw what appeared to be antennal duplications or other outgrowths (Fig. 1F-1G’, Supplemental Fig. S1). Some of these outgrowths resembled specific differentiated structures like antennae, maxillary palps, or even legs while others are difficult to classify (Fig. 1F-1G’, Supplemental Figure S1; quantification of relative “classified” versus “unclassified” outgrowths, Fig. 1K). On one occasion, we saw almost an entire leg growing from the eye with what may have been ommatidia on its distal tip (Supplemental Fig. S1C).

**Fig. 1:**
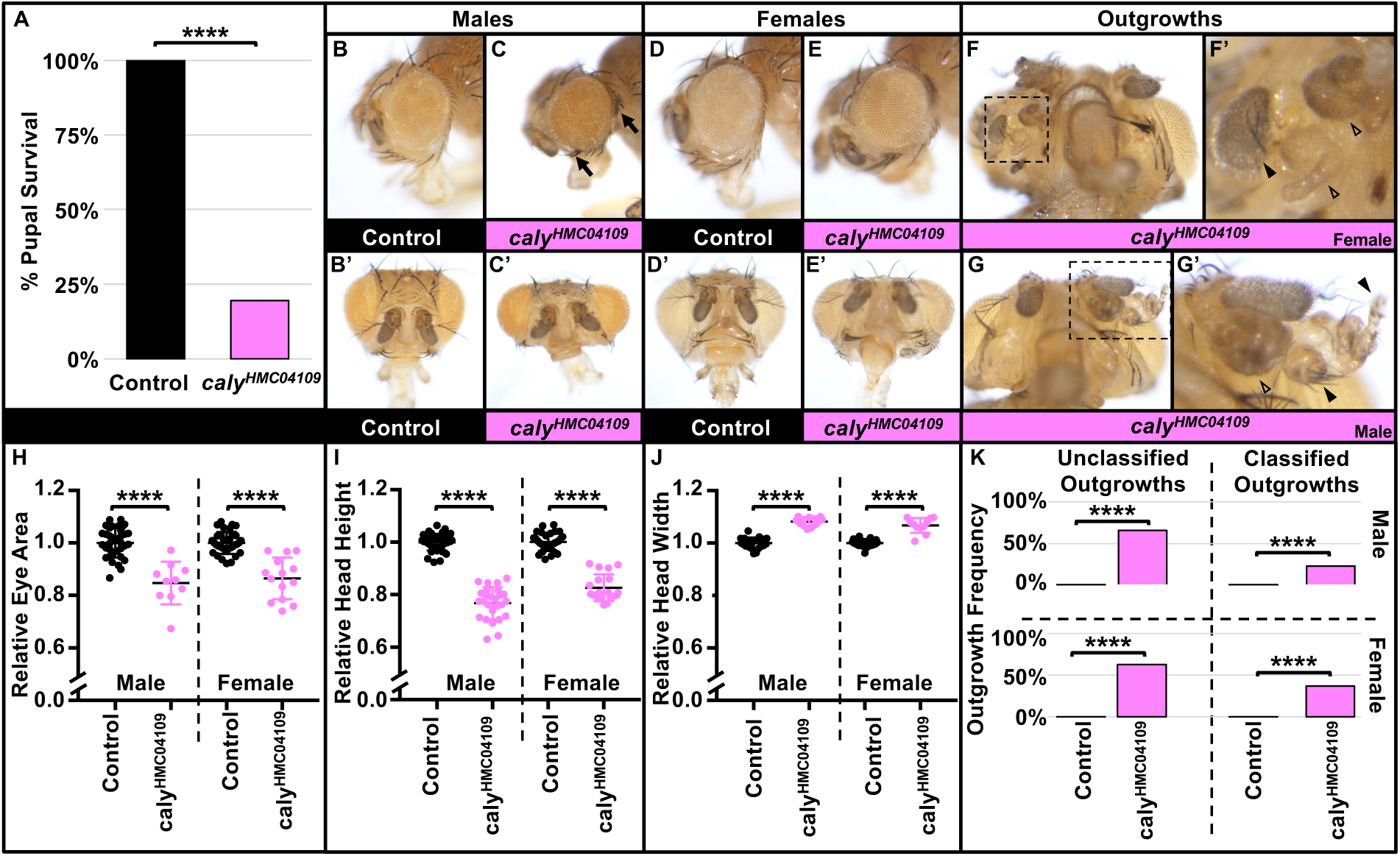
RNAi to *caly* using *ey-gal4* reduces pupal survival, causes eye shape and size phenotypes, and causes tissue outgrowths. Experiments were performed at 25°C. (A) Graph summarizing the percent pupal survival for control *ey-gal4/+* flies (left, black) versus driving RNAi to *caly* using *ey-gal4* and *caly^HMC04109^* (right, pink). RNAi to *caly* dramatically reduces survival of pupae to adulthood. **** indicates p<0.0001 in both Chi square and Fisher’s exact tests. (B-G’) eyes/heads in B-C’ and G-G’ show males and images in D-F’ show females. (B, D) *ey-gal4/+* eye profile showing control eye shape and size. (B’, D’) Anterior images of the heads in B,D. (C,E) RNAi to *caly* using *ey-gal4* (ey>*caly^HMC04109^*) causes a reduction in eye size, rounder eye shape, and change in bristle pattern (highlighted by the solid arrows in C). (C’, E’) Anterior view highlights that the head is shorter but wider. (F-G’) In addition to eye size and shape changes, eyes contain a variety of different outgrowths including outgrowths that appear to have differentiated into structures resembling antennae, maxillary palps, or legs (solid arrowheads) whereas others seem to lack obvious morphology associated with specific structures and cannot be classified morphologically based on visual inspection (arrowhead without fill). Additional view of outgrowths from the head in F-F’ is shown in Supplemental Fig. S1E-S1E’, and additional images of ey>*caly^HMC04109^* heads are shown in Supplemental Fig. S1, including an outgrowth of almost an entire leg from the eye in Supplemental Fig. S1C. (H-J) Graphs summarizing (H) relative eye area, (I) relative head height, and (J) relative head width for *ey-gal4/+* control males (first lane, black) and females (third lane, black) versus ey>*caly^HMC04109^* males (second lane, pink) and females (fourth lane, pink). **** indicates p<0.0001 in t tests. Eye image indicating how head height and width were measured is shown in Supplemental Fig. S2A. To highlight the head shape changes, we also graphed the ratio of width-to-height in Supplemental Fig. S2B. (K) Graphs quantifying the percent of eyes with outgrowths in *ey-gal4/+* controls (left lanes) versus ey>*caly^HMC04109^*(right lanes) when the outgrowths have unclassifiable outgrowths (left graphs) or outgrowths with classifiable morphology (such as legs, antennae, or palps) (right graphs) for males (upper graphs) and females (bottom graphs). **** indicates p<0.0001 in both Chi square and Fisher’s exact tests.

To establish if *ey>caly^HMC04109^* phenotypes resulted from knockdown of *caly* specifically and to rule out off-target effects, we generated eyes containing primarily *caly* mutant tissue for null allele *caly^2^* and catalytically inactive allele *caly^C131S^* by utilizing the FLP/FRT system and a cell-lethal mutation on the control 2R chromosome. *yweyFLP; FRT42D l(2)/40AFRT,FRT42D caly^2^* and *yweyFLP; FRT42D l(2)/40AFRT,FRT42D caly^C131S^* eyes contained “un-flipped” heterozygous tissue (red) and primarily *caly* mutant tissue (white) (Fig. 2C-D’, 2G-H’). These eyes were rough and showed reduced eye area, reduced head height, and increased head width phenotypes (Fig. 2I-2K) compared to control eyes (Fig. 2A-2A’, 2E-2E’, 2I-2K) as seen in *ey>caly^HMC04109^*eyes (Fig. 1, Supplemental Figs. S1-S2). These phenotypes also resembled eyes composed of primarily *Asx* mutant tissue (Fig. 2B-2B’, 2F-2F’). We also observe eye outgrowths (Fig. 2B, 2C).

**Fig. 2:**
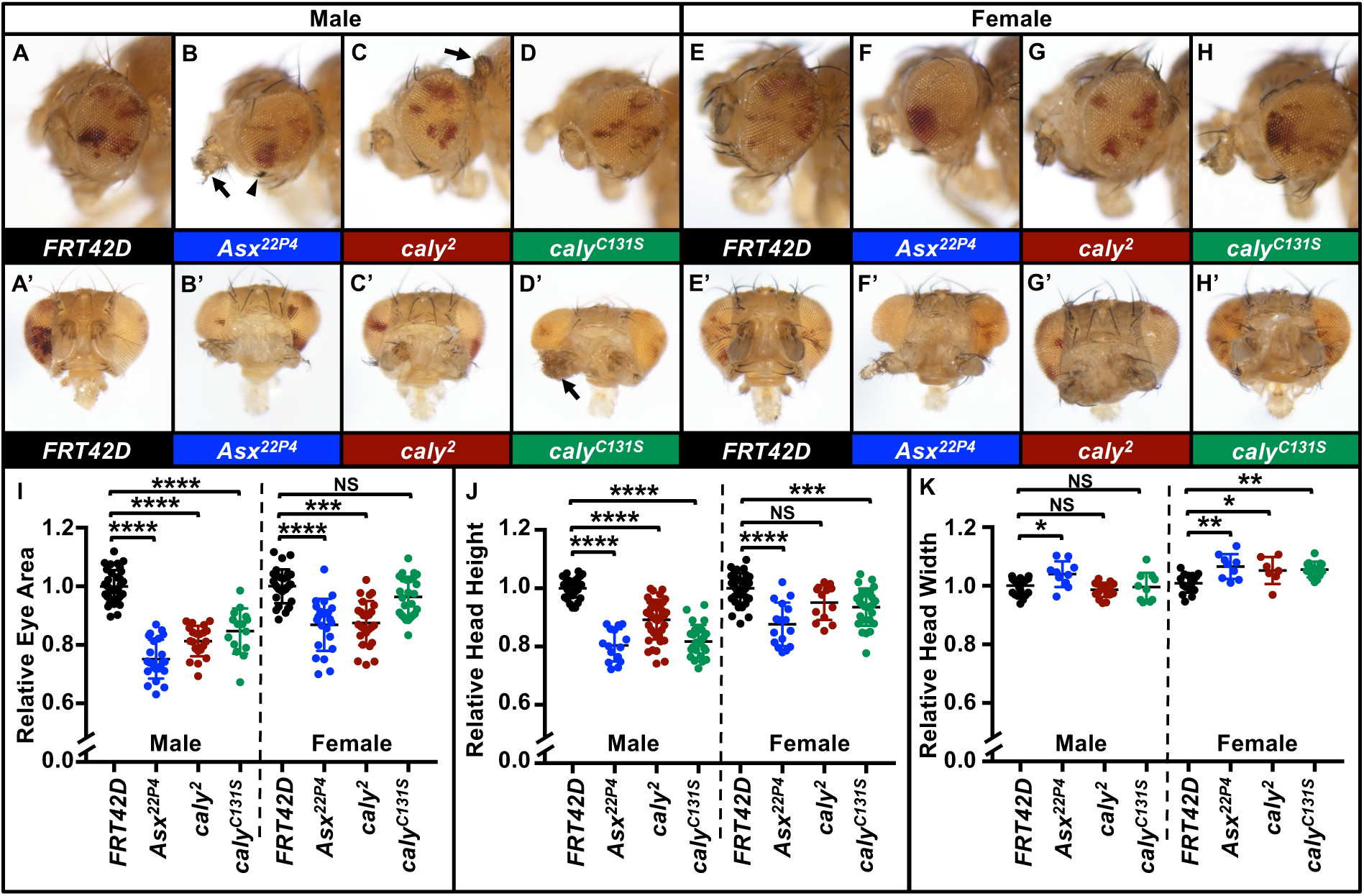
Eyes composed largely of homozygous mutant *caly* tissue in the eye phenocopies *caly* RNAi by changing eye shape and size and causing eye outgrowths. Experiments were performed at 25°C. Images in A-D’ show males and in E-H’ show females generated using *eyFLP,* an *FRT42D* chromosome with cell lethal mutation l(2)R11^1^ and a pW+ insertion, and an *FRT42D* chromosome either wild-type or containing mutations in *Asx* or *caly*. Due to the cell lethal mutation, “flipped” tissue homozygous for the l(2)R11^1^ chromosome dies. Remaining tissue is either red heterozygous “unflipped” tissue or white homozygous wildtype, *Asx*, or *caly* mutant tissue. Due to the lighting, white tissue in these images appears to have a yellow or orange hue. (A,E) Eyes containing *FRT42D* control tissue. (A’, E’) Anterior view showing eyes from A, E. (B, F) Eyes containing *Asx^22P4^* mutant tissue. (B’, F’) Anterior view showing eyes from B, F. (C, G) Eyes containing *caly^2^* null mutant tissue. (C’, G’) Anterior view showing eyes from C, G. (D, H) Eyes containing *caly^C131S^*catalytically inactive mutant tissue. (D’, H’) Anterior view showing eyes from D, H. Eyes containing *Asx* and *caly* mutant tissue are round and smaller with bristle abnormalities (example with an arrowhead in B) and with occasional outgrowths that differentiate into structures resembling antennal or other morphology (example with an arrow in B) or with no clear morphology (example with an arrow in D’). (I-K) Graphs summarizing (I) relative eye area, (J) relative head height, and (K) relative head width for eyes containing control tissue (lanes 1 and 5, black), *Asx^22P4^*mutant tissue (lanes 2 and 6, blue), *caly^2^* mutant tissue (lanes 3 and 7, dark red), and *caly^C131S^* mutant tissue (lanes 4 and 8, green). Eye image indicating how head height and width were measured is shown in Supplemental Fig. S2A. Males are shown in lanes 1-4 and females in lanes 5-8 for each graph. As with *caly* RNAi in Fig. 1, heads with *Asx* or *caly* mutant tissue have smaller eyes, reduced height, and increased width compared to controls. To highlight the head shape changes, we graphed the ratio of width-to-height in Supplemental Fig. S2C. In I-K, NS indicates not significant, * indicates p<0.05, ** indicates p<0.01, *** indicates p<0.001, and **** indicates p<0.0001.

We cannot rule out that off-target effects contribute to the phenotypes in *ey>caly^HMC04109^*flies, but the similarity of *ey>caly^HMC04109^* phenotypes compared to eyes containing primarily *caly^2^* or *caly^C131S^*mutant tissue would be consistent with these phenotypes resulting from loss-of-function in *caly*, not due to RNAi off-target effects on other genes. These phenotypes also resembled the phenotypes of dysregulating chromatin and interfering with Polycomb group proteins, their targets, and Pax6 [Zhu et al., 2018; Plaza et al., 2001; Arancio et al., 2010; Luque et al., 2007; Nègre et al., 2006], over-expression of OSA [Baig et al., 2010] or dysregulating other regulators of chromatin architecture like *Defective proventriculus* (*Dve*) [Puli et al., 2024]. In fact, dysregulation of many chromatin regulators and loss of mis-expression of Polycomb targets also demonstrate similar outgrowths, some of which appear to take on differentiated morphology resembling other structures such as cuticle or leg outgrowths we saw here [Plaza et al., 2001; Puli et al., 2024; Dong et al., 2002; Dey et al., 2009] including the ectopic legs seen upon overexpressing of *Antennapedia* (*Antp*) [Schneuwly et al., 1987] which is also consistent with the *Antp* misexpression seen in eye discs upon loss of *caly* [Brown et al., 2023].

### RNAi to *caly* in the differentiating eye causes mild overgrowth

*GMR-gal4* drives expression in cells posterior to the morphogenetic furrow in the differentiating eye [Freeman et al., 1996; Hay et al., 1994] and has also been described to express in the wing, midgut, salivary glands, and trachea [Escobedo et al., 2021; Li et al., 2012; Ray et al., 2015]. Inducing RNAi with *caly^HMC04109^* and *GMR-gal4* did not affect survival (Fig. 3A) and resulted in eyes of apparent normal morphology but increased eye area (quantified in Fig. 3B, eye examples Fig. 3D, 3F) compared to controls (Fig. 3C, 3E). The striking differences between eye phenotypes depending on driving RNAi with *ey-gal4* (Fig. 1) or with *GMR-gal4* might reflect different roles in early developing and actively proliferating tissue from roles in primarily differentiated tissue.

**Fig. 3:**
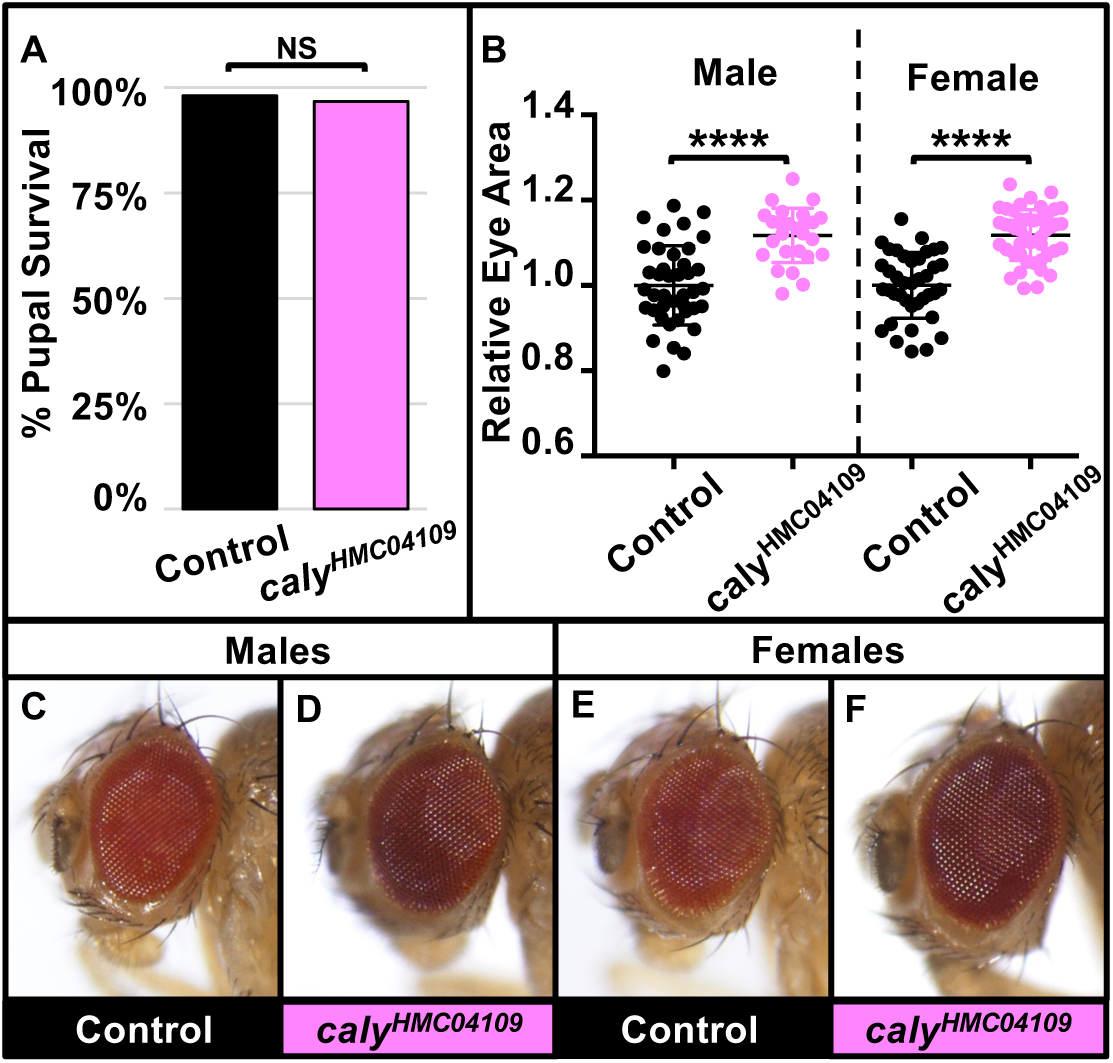
RNAi to *caly* using *GMR-gal4* does not affect morphology but causes increased eye size. Experiments in this figure were performed at 25°C. (A) Graph summarizing the percent pupal survival in control *GMR-gal4/+* flies (left, black) versus driving RNAi to *caly* using *GMR-gal4* and *caly^HMC04109^*(*GMR>caly^HMC04109^*) (right, pink). NS indicates not significant in both Chi square (p=0.087) and Fisher’s (p=0.12) exact tests. (B) Graph showing relative eye area in males (left two lanes) and females (right two lanes) of control GMR-gal4/+ flies (black, first and third lanes) and *GMR>caly^HMC04109^* flies (pink, second and fourth lanes) relative eye area males and females. Eye size increases by approximately 11.2%. The eye morphology appears normal despite the increased size. **** indicates p<0.0001 in t tests. (C,E) Control *GMR-gal4/+* eyes. (D,F) *GMR>caly^HMC04109^* eye.

### RNAi to *caly* in the dorsal wing caused wing vein abnormalities, shriveling, and “cupping.”

*ms1096-gal4* drives expression in the dorsal region of the wing disc pouch [Guillén et al., 1995; Rodan et al., 2002] but has also been described to drive expression in halteres, eye discs, and weak expression in ventral regions including the ventral cuticle [Jonchere et al., 2013; Shukla et al., 2014]. Inducing RNAi with *caly^HMC04109^* did not statistically significantly affect survival (quantified in Fig. 4A). *ms1096>caly^HMC04109^*wings showed a range of phenotypes (Fig. 4D-4D’’, 4G-4G’’, apparent size quantified in Fig. 4B, 4E) including disruption of wing vein pattern and loss of wing veins, blistering, crumpling/shriveling, a “cupped” wing shape, and a reduction in apparent overall wing size (crumpling and “cupping” interfered with measuring exact wing size) compared to controls (Fig. 4C, 4F). The wing vein abnormalities resemble those seen upon dysregulating other chromatin regulators and homeotic regulators such as HDAC complex components [Barnes et al., 2018], SWI-SNF chromatin remodeling complexes [Tian and Smith-Bolton, 2021], and trithorax group member *Absent small and homeotic 2* (*ash2*) [Amorós et al., 2002]. The shriveling/crumping phenotype resembled phenotypes seen for knocking down or mutating chromatin and homeotic genes including Asx [Bischoff et al., 2009], *DISCO Interacting Protein 1* (*DIP1*) [Bondos et al., 2004], *L(3)mbt* [Richter et al., 2011], *Antennapedia* (*Antp*) [Fang et al., 2022], and PcG protein *Pleiohomeotic* (*Pho*) [Harvey et al., 2013] and generally appear similar to what has been characterized as a “PcG syndrome” [de Ayala Alonso et al., 2007]. The “cupping” phenotype has also been seen upon reduction in HDAC proteins [Barnes et al. 2018] and *ash2* [Amorós et al., 2002], or deleting Polycomb response elements in Polycomb target genes [Sipos et al., 2007].

**Fig. 4:**
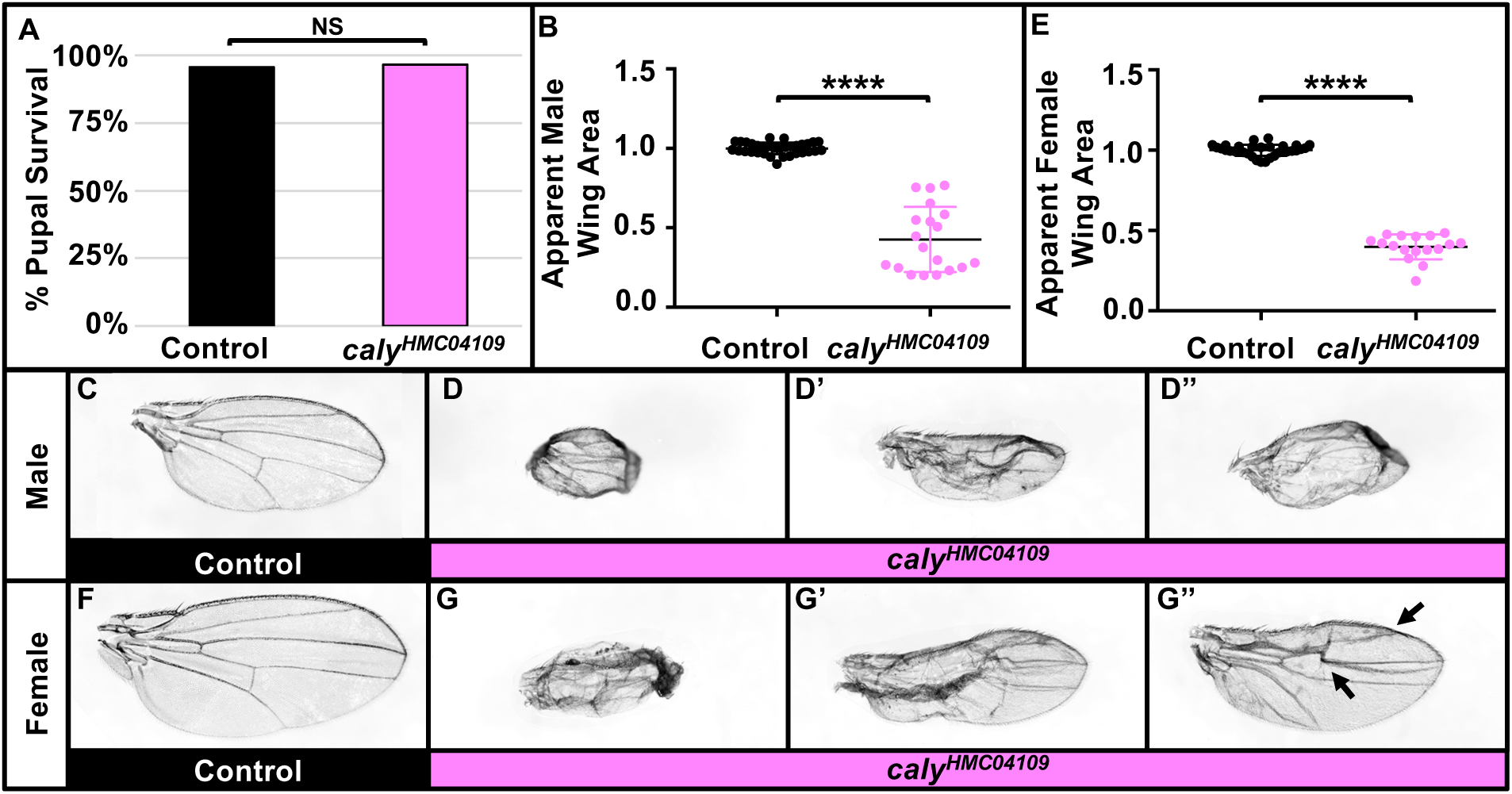
RNAi to *caly* using *ms1096-gal4* disrupts wing patterning and reduces wing size. Experiments in this figure were performed at 25°C. (A) Graph summarizing the percent pupal survival in control *ms1096-gal4/+* flies versus driving RNAi to *caly* using *ms1096-gal4* and *caly^HMC04109^* (*ms1096>caly^HMC04109^*). RNAi to *caly* does not affect pupal survival. NS indicates not significant in Chi Square (p=0.7468) and Fisher’s exact (p>0.9999) tests. (B-D’’) data for males; (E-G’’) data for females. (B, E) Relative apparent wing area decreases upon RNAi to *caly* (right, pink) compared to controls (left, black) for males (B) and females (E). (C, F) Control *ms1096-gal4* wing. (D-D’’, G-G’’) Both male (D-D’’) and female (G-G’’) *ms1096>caly^HMC04109^* wings have a range of phenotypes from very small wings with abnormal shapes including “cupping” of the wing or curling of the wing margins (D, G) to wings with moderate phenotypes that show less of a decrease in apparent wing area but still do not flatten properly even once inflated (D’, G’) to wings that with even milder phenotypes that still show abnormal patterning (arrows) (D’’, G’’).

### RNAi to *caly* across the wing caused wing vein abnormalities and curling at the wing margin

*c765-gal4* drives expression across the wing [Guillen et al., 1995; Nellen et al., 1996; de Celis et al., 1996] and also generally in in the thorax [Gomez-Skarmeta et al., 1996; Yang et al., 2012], in leg discs [Azpiazu and Morata, 2002], and in the brain [Rodan et al., 2002]. Inducing RNAi across the wing with *caly^HMC04109^*and *c765-gal4* resulted in a trend of reduced survival at 21°C (quantified in Fig. 5A) and this increased and became statistically significant at 25°C (quantified in Fig. 5F). The wings of flies that survived showed a number of phenotypic abnormalities including loss of wing vein material, a failure of some wing veins to reach the wing margin, buckling of tissue, and curling at the wing margin which was evident at 21°C (Fig. 5C, 5E) and increased at 25°C (Fig. 5H, 5J) compared to controls (Fig. 5B, 5D, 5G, 5I). These wing morphology phenotypes resembled but were weaker than those seen for *ms1096-gal4* and similar to those seen for other chromatin regulators described above.

**Fig. 5:**
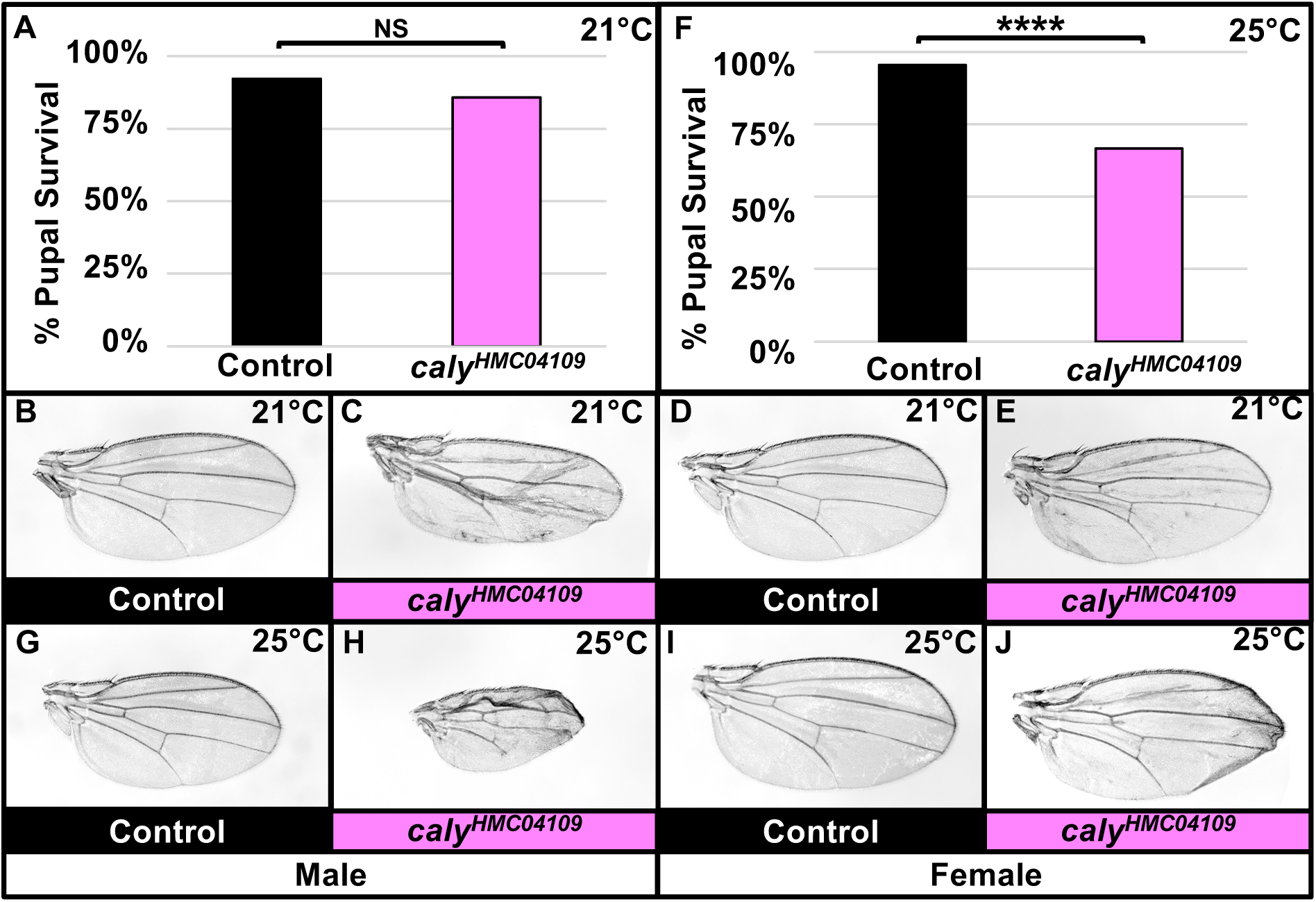
RNAi to *caly* across the wing using *c765-gal4* reduces survival and causes wing abnormalities. (A-E) Experiments in A-E were performed at 21°C. (A) Graph summarizing the percent pupal survival in control *c765-gal4/+* flies versus driving RNAi to *caly* using *c765-gal4* and *caly^HMC04109^*(*c765>caly^HMC04109^*). There is a reproducible trend of decreased survival upon *caly* RNAi, but this is not statistically significant. NS indicates not significant in Chi square (p=0.1788) and Fisher’s exact (p=0.2577) tests. (B,D) Control *c765-gal4/+* wing. (C,E) c765>*caly^HMC04109^* wing. c765>*caly^HMC04109^* wings have disrupted wing patterning including loss of anterior and posterior crossveins. Male wings are shown in B-C, and female wings in D-E. (F-J) Experiments in F-J were performed at 25°C. (F) Graph summarizing the percent pupal survival in control *c765-gal4/+* flies versus c765>*caly^HMC04109^* flies. Reproducibly, there is statistically significant decreased survival upon *caly* RNAi at the higher temperature of 25°C compared to the survival at 21°C shown in A. **** indicates p<0.0001 in both Chi square and Fisher’s exact tests. (G,I) *c765-gal4/+* wing. (H,J) c765>*caly^HMC04109^* wing. RNAi to *caly* disrupts wing patterning and causes curling at the wing margin, and this is more obvious in male wings. Male wings are shown in G-H, and female wings in I-J.

The lethality of driving *caly* RNAi with drivers whose primary expression pattern is in tissues that themselves are not required for viability (e.g., eyes, wings) raises interesting questions regarding the cause of this lethality. It is possible *caly* knockdown in these tissues caused secretion of factors or other systemic effects. Another possibility is that lethality resulted from effects on other tissues where these Gal4 drivers also induce expression. As reviewed earlier, BAP1 is a known tumor suppressor and metastasis suppressor, and we noted tumor-like outgrowths in the eye. Therefore, the lethality of *caly* RNAi in certain tissues could also reflect metastasis of cells to other sites. In fact, BAP1 loss is associated with worse prognosis in UM. Pursuing these possible mechanisms in future work may shed light on the requirement for *caly/BAP1* in development with implications for disease.

In addition to a role in cancer, heterozygous mutations in *BAP1* are implicated in a neurodevelopmental syndrome known as Küry-Isidor syndrome (KURIS). This is characterized by developmental delay that affects walking and speech [Küry et al., 2022]. Although we cannot rule out off-target effects resulting from RNAi allele *caly^HMC04109^* in the differentiating eye (Fig. 3) or the wing (Figs. 4-5), eyes containing primarily *caly^2^* and *caly^C131S^* (Fig. 2) phenocopied driving *caly^HMC04109^* to induce RNAi in the early eye (Fig. 1, Supplemental Fig. S1) as well as phenotypes associated with loss of other chromatin regulators. This is consistent with the *caly^HMC04109^* allele being a useful tool to assess further the mechanisms underlying the role of BAP1 defects in cancer and in Küry-Isidor syndrome. Importantly, some eye outgrowths upon *caly* loss using both RNAi (Fig. 1, Supplemental Fig. S1) and mutant alleles (Fig. 2) resemble tumor-like growths seen for other cancer models [Yan et al., 2009; Pagliarini and Xu, 2003; Uhlirova et al., 2005; Brumby and Richardson, 2003; Ho et al., 2015] while others resemble homeotic transformations. Therefore, recapitulating *caly* loss in the early eye using *caly^HMC04109^*, *caly^2^*, and *caly^C131S^* could make an excellent developmental context to study the relationship between epigenetic dysregulation and tumor initiation to be pursued in future work.

## MATERIALS AND METHODS

### Rigor and Reproducibility

The reported work represents reproducible experiments that reflect a minimum of three well-controlled, independent trials. For most experiments, at least one set of trials was done by a different lab member than the other two sets of trials to reduce the chance of replicating unintended observer bias.

### Statistical analysis

Eye area, head height, head width, and wing area were measured with ImageJ software. Raw measurements in pixels and lethality data were normalized and graphed using Excel (Fig. 1A, 1K, 3A, 4A, 5A, 5F) and GraphPad Prism (Fig. 1H-1J, 2I-2K, 3B, 4B, 4E). Categorical analysis to analyze survival (Fig. 1A, 3A, 4A, 5A, 5F) or type of outgrowths (Fig. 1K) used Chi square and Fisher exact tests as appropriate calculated in GraphPad Prism. T-test (Fig. 1H-1J, 3B, 4D, 4E) and one-way ANOVA analysis with multiple comparisons (Fig. 2I-2K) assessed changes in eye area, head height, head width, or wing size. P values, raw measurements, and normalized values are listed in Supplemental File S1.

### *Drosophila* experiments

Crosses at the indicated temperatures were set up on standard *Drosophila* medium as in our previous work [Yan et al. 2009; Yan et al., 2010, Washington et al., 2020; Reimels et al., 2024]. In each trial for each experiment, crosses used food prepared in the same batch and were incubated in close proximity to experience the same environment to rule out unintended environmental variables or food batch variations. Gal4 drivers were obtained from the Bloomington *Drosophila* Stock center or other labs in the *Drosophila* community (for details, please see Table 1). *caly* RNAi caly^HMC04109^, and cell lethal *l(2)cl-R11^1^* were from the Bloomington Stock center, and *caly* and *Asx* alleles were graciously provided by Dr. J. Müller [de Ayala Alonso et al., 2007].

**Table 1:** Table of reagents used with corresponding identifiers.

| ***Drosophila* Strains** |  |  |
| --- | --- | --- |
| **Strain** | **Source** | **Identifier** |
| *w*<sup>1118</sup> | The fly community and Bloomington *Drosophila* Stock Center (BDSC) | BL-3605, BL-5905 and others<br>RRID:BDSC_3605, RRID:BDSC_5905 |
| *caly*<sup>HMC04109</sup> | BDSC | Can be obtained from BDSC, 56888<br>RRID:BDSC_56888 |
| *ey-gal4* | Hariharan lab |  |
| *c765-gal4* | BDSC, NYC fly community | Can be obtained from BDSC, BL-36523<br>RRID:BDSC_36523 |
| *ms1096-gal4* | BDSC | Can be obtained from BDSC, BL-8696<br>RRID:BDSC_8696 |
| *GMR-gal4* | BDSC | Can be obtained from BDSC, BL-8605<br>RRID:BDSC_8605 |
| *FRT42D* | BDSC | Can be obtained from BDSC, BL-5626<br>RRID:BDSC_5626 |
| *y, w, eyFLP, GMR-lacZ; FRT42D, l(2)cl-R11<sup>1</sup>, w+/CyO* | BDSC | Can be obtained from BDSC, BL-5617<br>RRID:BDSC_5617 |
| *FRT40, FRT42D y+ caly<sup>2</sup> /CyO,ubi-GFP* | Gift from Müller Lab |  |
| *FRT40, FRT42D y+ caly<sup>C131S</sup> /CyO,ubi-GFP* | Gift from Müller Lab |  |
| *FRT40, FRT42D y+*<br>*Asx<sup>C131S</sup>/CyO,ubi-GFP* | Gift from Müller<br>Lab |  |
| **Software** |  |  |
| **Program** | **Source** |  |
| Image J | https://imagej.net/ij/ |  |
| GraphPad Prism | https://www.graphpad.com/scientific-software/prism/ |  |
| Microsoft Excel | https://www.microsoft.com/Microsoft/Excel/ |  |

### Pupal lethality/survival experiments

For survival experiments, pupal cases of the indicated genotypes were scored as dead (in which dead pupae remained in the pupal cases) or empty (from which surviving flies had eclosed) and counted at 26 days (21°C) or 18 days (25°C) as in our previous study [Singh et al., 2023]. Dead pupal cases are easily distinguished from empty pupal cases or from developing pupae that are still alive and have not yet eclosed.

### Image analysis and processing

Adult eyes and wings were photographed using a Nikon DS-Fi3 microscope camera and saved as TIFF files. Eye images were adjusted for clarity; wing images were converted to grayscale and adjusted using Adobe Photoshop. Brightness and contrast adjustments were applied to the entire images. Genotypes are summarized below and identifiers are listed in Table 1.

### Genotypes of flies in images or graphs

*w; ey-gal4/+* (Fig. 1B-1B’, 1D-1D’; left genotype in graphs in Fig. 1A, 1H-1K, Supplemental Figure S2B)

*w; ey-gal4/caly^HMC04019^* (Fig. 1C-1C’, 1E-1E’, 1F-1G’, Supplemental Fig. S1A-S1E’; right genotype in graphs in Fig. 1A, 1H-1K, Supplemental Fig. S2B)

*y w eyFLP; FRT42D/FRT42D l(2)cl-R11^1^* (Fig. 2A-2A’, 2E-2E’; left-most genotype in graphs in 2I-2K, Supplemental Fig. S2C)

*y w eyFLP; FRT42D Asx^22P4^/FRT42D l(2)cl-R11^1^*(Fig. 2B-2B’, 2F-2F’; second genotype in graphs in 2I-2K, Supplemental Fig. S2C)

*y w eyFLP; FRT42D caly^2^/FRT42D l(2)cl-R11^1^*(Fig. 2C-2C’, 2G-2G’; third genotype in graphs in 2I-2K, Supplemental Fig. S2C)

*y w eyFLP; FRT42D caly^C131S^/FRT42D l(2)cl-R11^1^* (Fig. 2D-2D’, 2H-2H’; right-most genotype in graphs in 2I-2K, Supplemental Fig. S2C)

*w; GMR-gal4/+* (Fig. 3C, 3E; left genotype in graphs in Fig. 3A-3B)

*w; GMR-gal4/caly^HMC04019^* (Fig. 3D, 3F; right genotype in graphs in Fig. 3A-3B)

*w, ms1096-gal4* (Fig. 4C, left genotype in graph in Fig. 4A, 4B)

*w, ms1096-gal4/+* (Fig. 4F, left genotype in graph in Fig. 4A, 4E)

*ms0196-gal4; caly^HMC04019^/+* (Fig. 4D-4D’’; right genotype in graph in 4A, 4B)

*ms0196-gal4/+; caly^HMC04019^/+* (Fig. 4G-4G’’; right genotype in graph in 4A, 4E)

*w; c765-gal4/+* (Fig. 5B, 5D, 5G, 5I; left bar in graphs in Fig. 5A, 5F)

*w; caly^HMC04019^/+; c765-gal4/+* (Fig. 5C, 5E, 5H, 5J; right bar in graphs in 5A, 5F)

## DATA AVAILABILITY STATEMENT

*Drosophila* strains used in this work (listed in Table 1) have been published previously [de Ayala Alonso et al., 2007] or available from public stock centers. Raw data, normalized data for graphs in Figs. 1-5, and p values are listed in Supplemental File S1. The authors affirm that all data necessary for interpreting the data and drawing conclusions are present within the article text, the figures, table, and Supplemental File S1.

## Supporting information

Luf et al. Supplemental File S1

## ACKNOWLEDGMENTS

We thank M Mlodzik, U Weber, TK Das, J Chipuk, P Rangan, ZQ Pan, and the New York Fly community. We thank D. Sethi, P. Karunaraj, K. Kalafsky, K. Braden, and F. Rosemann for assistance. We thank Dr. J. Müller and his lab for generously providing us with fly stocks. We thank the Bloomington Drosophila Stock Center (NIH P40OD018537) for providing fly stocks and Flybase (NIH 5U41HG000739) for access to sequence information.

## Supplemental Figures

**Supplemental Figure S1:**
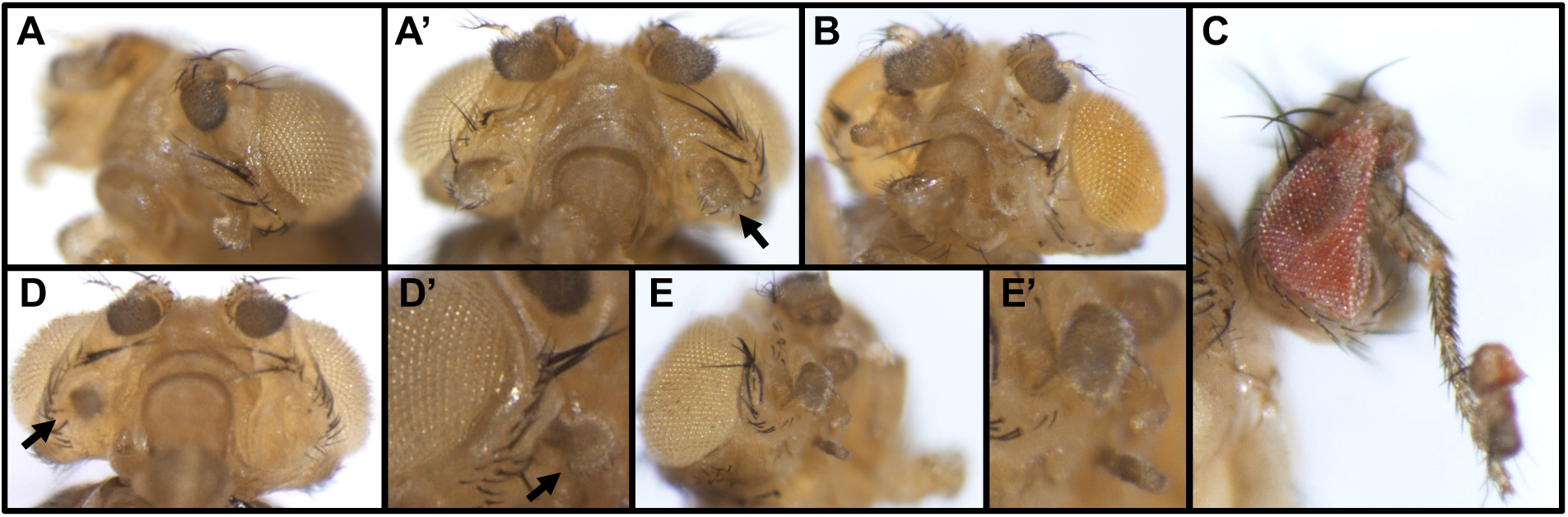
Additional images of ey>*caly^HMC04109^* heads highlighting outgrowths. (A-E) Additional examples of ey>*caly^HMC04109^* heads showcasing outgrowths (solid arrows) and bristle abnormalities. (A-A’) Alternate views of the same head showing outgrowths on both sides. (B) Another example of a head with an outgrowth where the outgrowth lacks obvious morphology and is hard to classify. (C) Example of a case where almost an entire leg is growing out of the head with additional tissue on the tip including ommatidia. We have seen multiple cases where outgrowths resemble leg segments, however, this is the only case we have seen where the outgrowth is this long. (D-D’) Alternate views of another head, with an enlarged view in D’. (E-E’) Alternate views of head shown in Fig. 1F-1F’ (enlarged in E’) highlighting the outgrowths. Females are shown in A-A’, C-E, and a male is shown in B.

**Supplemental Figure S2:**
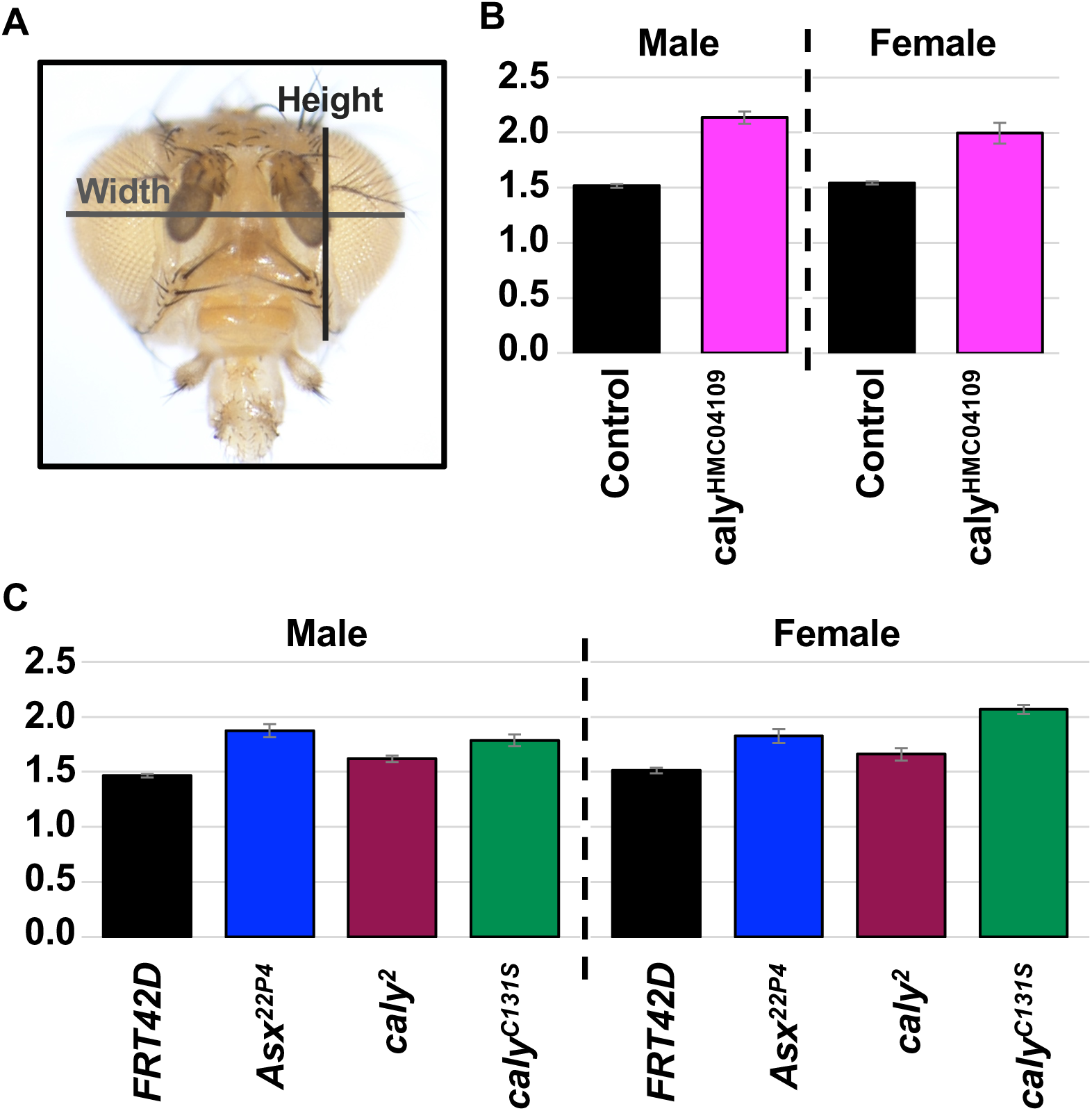
Decreased *caly “*flattens*”* heads as shown by increased ratio of width-to-height. in. (A) Diagram indicating how head height and width were measured for graphs in Fig. 1I-1J and Fig. 2J-2K. Anterior image of a head with lines overlaid. Vertical line indicates how head height measurements were taken for graphs in Fig. 1I and Fig. 2J and horizontal line indicates how head width measurements were taken for graphs in Fig. 1J and Fig. 2K. (B) Graph indicating the Ratio of width-to-height for ey>*caly^HMC04109^* (pink, second and fourth lanes) versus *ey-gal4/+* (black, first and third lanes) heads for males (lanes one and two) and females (lanes three and four). (C) Graph indicating the Ratio of width-to-height for heads containing largely *FRT42D* control tissues (black bars, first and fifth lanes), *Asx^22P4^*mutant tissues (blue, second and sixth lanes), *caly^2^* mutant tissue (dark red, third and seventh lanes), and *caly^C131S^* mutant tissues (fourth and eighth lanes) for males (lanes one through four) and females (lanes five through eight). Error bars in B-C reflect additive relative standard error of the mean (SEM) from individual height and width calculations. SEM for the height and width for graphs in Figs. 1 and 2 was calculated in GraphPad Prism and divided by the relevant means to indicate relative SEM. Relative SEM for height was added to relative SEM for width and then applied to the ratio of width to height. Statistical analysis for individual height and width measurements is shown in the main figures and summarized in Supplemental File S1.

